# Vaginal bacteria elicit acute inflammatory response in fallopian tube organoids: a model for pelvic inflammatory disease

**DOI:** 10.1101/2023.02.06.527402

**Authors:** Bo Yu, Edward B. Nguyen, Stephen McCartney, Susan Strenk, Daniel Valint, Congzhou Liu, Catherine Haggerty, David N. Fredricks

## Abstract

**Objective:** To facilitate *in vitro* mechanistic studies in pelvic inflammatory disease (PID) and subsequent tubal factor infertility, as well as ovarian carcinogenesis, we sought to establish patient tissue derived fallopian tube (FT) organoids and to study their inflammatory response to acute vaginal bacterial infection.

**Design:** Experimental study.

**Setting:** Academic medical and research center.

**Patients:** FT tissues were obtained from four patients after salpingectomy for benign gynecological diseases.

**Interventions:** We introduced acute infection in the FT organoid culture system by inoculating the organoid culture media with two common vaginal bacterial species, *Lactobacillus crispatus* and *Fannyhessea vaginae*.

**Main Outcome Measures:** The inflammatory response elicited in the organoids after acute bacterial infection was analyzed by the expression profile of 249 inflammatory genes.

**Results:** Compared to the negative controls that were not cultured with any bacteria, the organoids cultured with either bacterial species showed multiple differentially expressed inflammatory genes. Marked differences were noted between the *Lactobacillus crispatus* infected organoids and those infected by *Fannyhessea vaginae*. Genes from the C-X-C motif chemokine ligand (CXCL) family were highly upregulated in *F. vaginae* infected organoids. Flow cytometry showed that immune cells quickly disappeared during the organoid culture, indicating the inflammatory response observed with bacterial culture was generated by the epithelial cells in the organoids.

**Conclusion:** Patient tissue derived FT organoids respond to acute bacterial infection with upregulation of inflammatory genes specific to different vaginal bacterial species. FT organoids is a useful model system to study the host-pathogen interaction during bacterial infection which may facilitate mechanistic investigations in PID and its contribution to tubal factor infertility and ovarian carcinogensis.

## INTRODUCTION

The fallopian tube (FT) is responsible for gamete and early embryo transportation, and is the site of fertilization and preimplantation embryonic development in the first 3 to 4 days of human life. The pathological significance of the FT includes: 1) As the conduit between the peritoneal and uterine cavity, FT is a site of pelvic inflammatory disease (PID) in which ascending vaginal bacterial infections affect the uterus, FT and ovaries. Untreated PID can lead to infertility due to tubal damage and occlusion, as well as tubal ectopic pregnancy, and repeated PID is a risk factor for developing ovarian cancer^1^. 2) FT is the site of origin for a large percentage of epithelial ovarian cancers as shown in studies of human precancerous lesions and in mouse models^2–8^. Based on consistent evidence of decreased ovarian cancer risks after bilateral tubal ligation^9–11^, we postulate that ascending bacterial infection and resulting inflammation of FT could be a contributing factor in ovarian carcinogenesis. 3) Endometriosis, which is a chronic inflammatory disease involving ectopic endometriotic lesions, is another risk factor for ovarian cancer^12^. FT not only is one of the sites of endometriosis, but also may contribute to the pathogenesis of endometriosis through retrograde menstruation and the spread of ascending pathogens and inflammatory substances^13^.

Establishing an *in vitro* model system for FT infection will facilitate mechanistic studies in PID and subsequent tubal factor infertility, as well as ovarian carcinogenesis. Organoids have been shown to preserve the original tissue characteristics in many different types of tissues and have emerged to serve as *in vitro* models for many diseases^14^. FT epithelial organoids have been derived from FT tissues^15^ or differentiated from induced pluripotent stem cells^16^, in which the secretory and ciliated cells are self organized into a single layer of columnar epithelium surrounding a lumen, mimicking the FT epithelium (FTE). Several studies have been published on FT organoids^15–18^ but few have utilized FT organoids to study infectious diseases, although one study with the emphasis on cancer transformation did establish chronic infection of FT organoids with *Chlamydia* and showed a molecular phenotype associated with aging in epithelial cells after prolonged culture^17^.

We sought to establish a FT organoid infection model to study common vaginal bacteria and their interaction with the FT epithelial cells in the organoids. The reasons for choosing vaginal bacteria to study include: 1) In our previous studies, we have shown that vaginal bacteria can ascend into the uterus and become part of the uterine microbiome^19^. 2) Recent studies demonstrated that vaginal bacteria were identified in the FT in women without overt signs of infection^20,21^. 3) Some vaginal bacterial species and genera that are associated with bacterial vaginosis (a common vaginal infection caused by bacterial overgrowth of anaerobes) including *Fannyhessea vaginae*, *Sneathia*, and *Megasphaera*, have been isolated from the upper genital tract of people with PID including the FT^22–25^. Taken together, this evidence indicates potential physiological or pathological significance of vaginal bacterial species in the upper female genital tract including the FT.

Two common vaginal bacterial species were chosen for study: *Lactobacillus crispatus* (*L. crispatus*) which represents part of the healthy vaginal microbiota, and *Fannyhessea vaginae (F. vaginae)*, which is a pathogen linked to PID^23^ and bacterial vaginosis^26^. In this pilot study, we introduced acute infection using non-pathogenic *L. crispatus* and pathogenic *F. vaginae* and studied the inflammatory response of the FT organoids, demonstrating the feasibility of establishing an infectious disease model in the FT organoids and the divergence of some of the host genes whose expression is influenced by these two bacterial species.

## METHODS

### Tissue collection and fallopian tube organoid culture

The study was approved by the institutional review board at University of Washington. We obtained informed written consents from each patient before her surgical procedure. FT tissues from 4 patients were collected in the operating room during salpingectomy for benign gynecological indications. The FT tissues were minced and digested with collagenase type I (Sigma) for one hour at 37°C. The dissociated mucosal cells were seeded and cultured to organoids in Matrigel following the published protocols with modifications^15^. Briefly, the cells were cultured in 2D culture media with progenitor growth factors as previously published^15^ at 37°C, 5% CO2 in a humidified incubator until 70-80% confluent. The epithelial cells were then detached and seeded in 75-80% Matrigel (Corning). The Matrigel domes were overlaid with 3D expansion medium as previously described^15^ and incubated at 37°C, 5% CO2 in a humidified incubator until the time for passage which on average takes 14-21 days. For the passage of organoids, after the Matrigel was dissolved with cold PBS, mechanical fragmentation was achieved by vigorous pipetting. Sheared organoids were centrifuged and resuspended in cold Matrigel and replated at the desired density. Organoid reformation were usually observed within 2-3 days after the passage. The culture medium was exchanged every 3–4 days.

### Immunohistochemistry (IHC)

IHC was perfomed as described previously with modifications^18^. FT organoids were released from the Matrigel with cold 4% paraformaldehyde (Fisher Scientific) and fixed for 2 hours before being embedded in Specimen Processing HistoGel™ (Thermo Fisher). The organoids encapsulated in the Histogel were dehydrated in an ascending series of alcohol, followed by isopropanol and acetone at room temperature. The dehydrated organoids were paraffin-embedded and cut into 5 μm sections on a microtome. Hematoxylin & Eosin (H&E) staining and IHC were carried out following standard procedures. The primary antibodies used in IHC included: Mouse anti-E Cadherin (BD, 610181); Rabbit anti-Pax8 (ProteinTech, 10336-1-AP); Mouse anti-acetylated Tubulin (Sigma, T7451).

### Bacterial preparation

*Lactobacillus crispatus* DNF00082 and *Fannyhessea vaginae* DNF00720 were inoculated from frozen stock onto Brucella agar supplemented with 5% laked sheep blood, hemin and Vitamin K (BRU) (Hardy Diagnostics, Santa Maria, CA) and grown at 37°C in anaerobic conditions (90% N2, 5% H2, 5% CO2). Isolates were subcultured twice on BRU or in modified peptone-yeast-glucose medium supplemented with 1% yeast extract and 1% glucose (PYG-mod-YG). A final culture for each isolate of 20mL PYG-mod-YG was centrifuged and pelleted and the pellets were reconstituted with 400 to 500 μL of PYG-mod-YG. Serial dilutions were made from 10 μL of reconstituted pellet and plated onto BRU to count colony forming units (CFU). The remaining reconstituted pellet was frozen with glycerol to make a 10% glycerol stock solution. CFU were counted once colonies had grown. Based on the final CFU count, portions of the 10% glycerol stock solutions were diluted with PYG-mod-YG to make 12 μL aliquots of 10^6^ CFU each. The aliquots were kept frozen at −80°C until they were inoculated into the organoids. Uninoculated aliquots of 12 μL PYG-mod-YG were also frozen with the isolate aliquots as negative controls. The 10% glycerol stock solutions were plated again for CFU counts on the same day as organoid inoculation to verify isolate viability after freezing.

### Co-culture of bacteria and organoids

The FT organoid culture media was changed to antibiotics-free media 24 hours before the acute infection experiment, and the media remained antibiotics-free during the following experiment. The FT organoids from each patient (n=4 patients) was inoculated with 10^6^ CFU of *Lactobacillus cripspatus* or *Fannyhessea vaginae* per well (n=3 replicates for each bacteria strain for 3 patients, n=1 for each strain for the first patient). Three negative controls (n=3 replicates for each control) included: PBS inoculation, sterile bacterial suspension media innoculation, and organoids alone without any innoculation. After 24 hours of incubation, the organoids were released from Matrigel, and dissociated into single cells as described below before RNA extraction.

### RNA extraction and gene expression profiling

Total RNA was extracted using Trizole (Life Technologies) according to the supplier’s protocol. The RNA quality was assessed using Tapestation and the RNA quantity was measured with Qubit fluorometer. All samples had RNA integrity number (RIN) > 9.0. Each sample was run on a lane of nCounter Inflammation Panel (Nanostring). The data was analyzed using the Nanostring nSolver analysis software 4.0 following the manufacturer’s instructions.

### Flow cytometry

After being released from the Matrigel with cold PBS, the FT organoids were dissociated into single cells with dispase (at 37°C for 30 minutes) and 0.25% trypsin (at 37°C for 2 minutes) digestions. After viability check, the cell suspension was filtered through 40 μm filter to get single cells for flow cytometry. Control samples were obtained from previously collected peripheral blood mononuclear cells (PBMC) which were isolated by Ficoll Histopaque (Pharmacia Biotech) gradient centrifugation at a density of 1.077 g/mL and cryopreserved in dimethylsulfoxide (DMSO). Dissociated organoids and control PBMCs were labeled with a fluorescent marker for viability (LIVE/DEAD Violet, Invitrogen) and anti-human CD45 (FITC conjugated, BD Biosciences cat#555482). The cells were fixed (Fix/Perm Buffer, BD Biosciences) and analyzed by flow cytometry using a FACS Canto II. The frequency of the CD45+ population was determined by percentage of total live cells (FlowJo).

## RESULTS

From FT tissues collected from consented patients at the time of surgery, we have established robust FT organoid cultures which can be repeatedly passaged for months while maintaining the same morphological and histological appearance (Figure 1). Hematoxylin and eosin (H&E) staining and immunohistochemistry demonstrated luminal structures with single-layer columnar epithelium consisting of ciliated and secretory cells (positive for Tubulin and PAX8 cell markers, respectively) (Figure 1). The ciliated cells are apical with the cilia facing the lumen, away from the basal membrane (positive for E-cadherin) (Figure 1).

**Figure 1.**
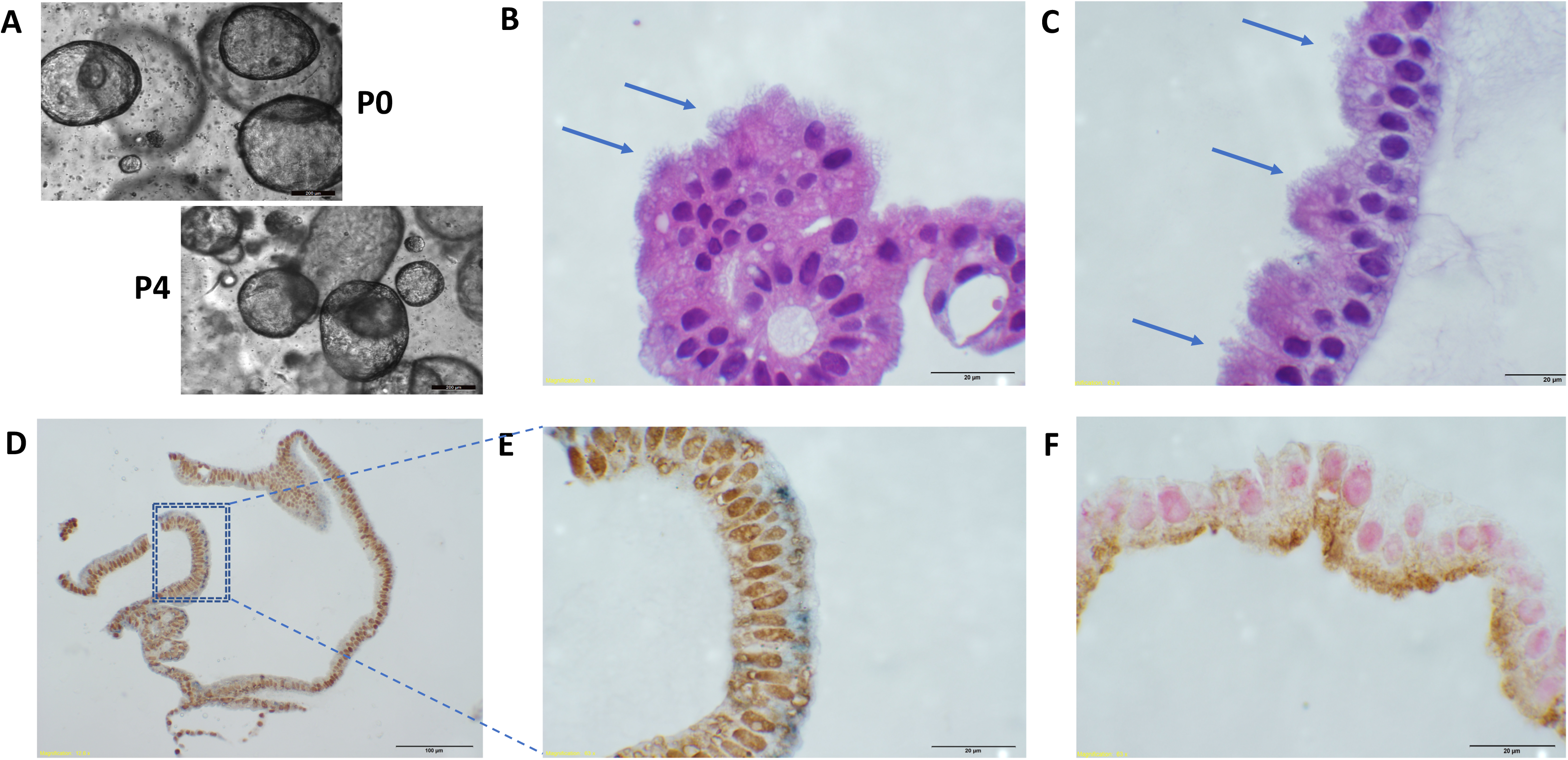
Characteristic appearance of Fallopian tube organoids. A. Morphology remains the same after 4 passages. B-C. H&E staining showing columnar epithelium with cilia (arrows) facing towards the lumen. D-E. IHC with PAX8 (brown) labeling secretory cells, tubulin (blue) labeling cilia. Dashed box in D is shown in higher magnification in E. F. IHC with PAX8 (pink) labeling secretory cells, E-cadherin labeling basal membrane. Scale bars: A: 200 μm; D: 100 μm; others: 20 μm.

We introduced acute infection in the FT organoid culture system by inoculating the organoid culture media with two common vaginal bacterial species, *Lactobacillus crispatus* (*L. crispatus*) and *Fannyhessea vaginae (F. vaginae)*. After 24 hours of culturing with or without one of the two bacterial species, no notable morphological changes were observed among the three groups (Figure 2). RNA was extracted from the organoids and the human gene expression profiles were analyzed using Nanostring Inflammatory panels which quantified the expression levels of 249 genes involved in the inflammatory process (Figure 2).

**Figure 2.**
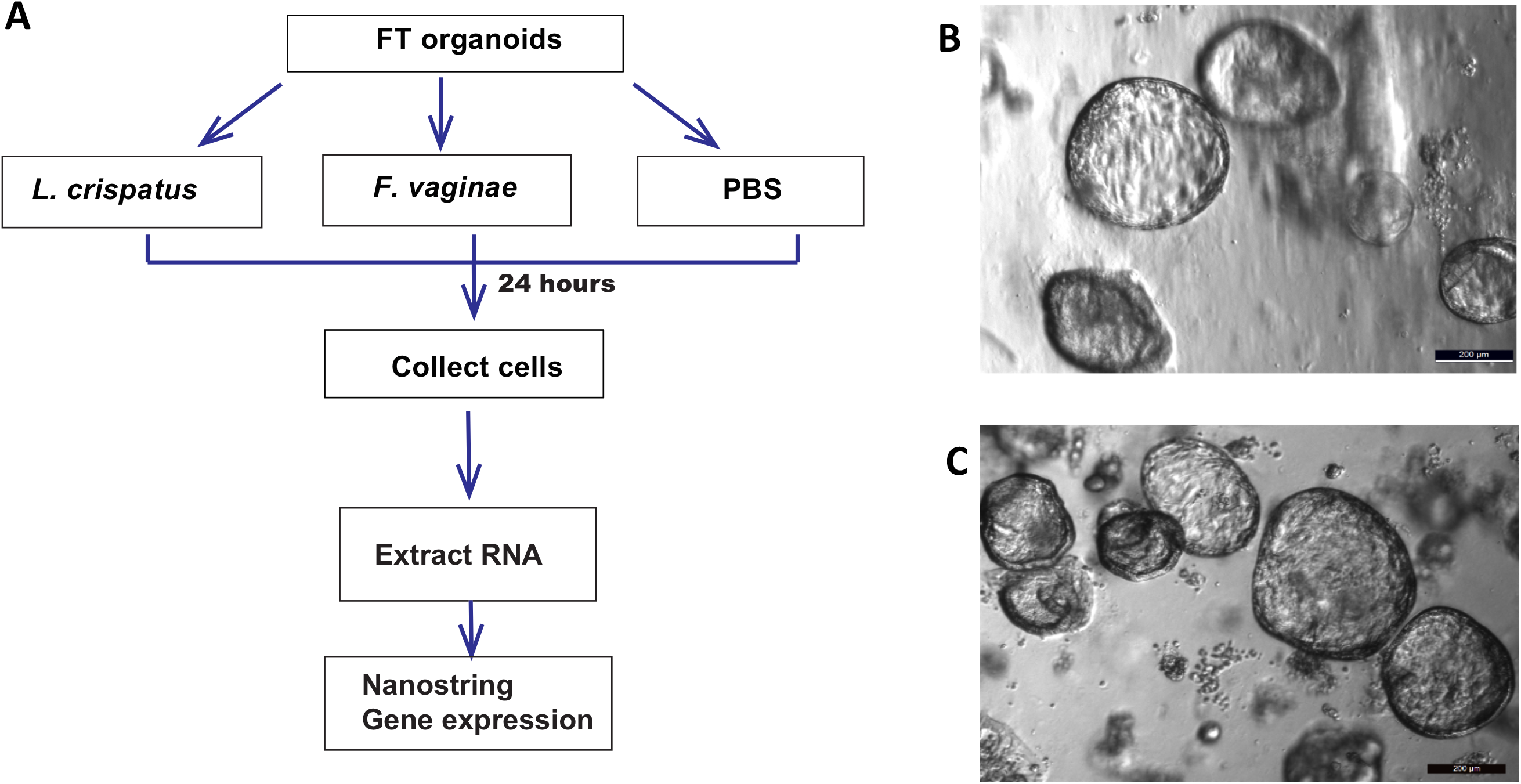
Experimental design and representative samples. A. Experimental design. B-C. Morphology of FT organoids remained unchanged before (B) and after (C) 24 hours of bacterial culture.

Among the 249 genes analyzed, 162 genes were expressed by all organoid samples and 71 genes were not expressed by any organoid sample. Organoids from each individual showed different inflammatory responses, however, the differential expression pattern between the experimental groups and in comparison with the controls can be clearly observed regardless of the inter-individual heterogeneity (Figure 3). Compared to the negative controls that were not cultured with any bacteria, the organoids cultured with either bacterial species showed multiple differentially expressed inflammatory genes (Figure 4), indicating the epithelial cells in the organoids can respond to acute bacterial infection and generate inflammatory responses. Notably, multiple members of the CC-chemokine ligand (CCL) family were upregulated in both *F. vaginae* and *L. crispatus* infected organoids, while multiple genes in the mitogen-activated protein kinase (MAPK) family are downregulated in *L. crispatus* infected organoids (Figure 4). When the gene expression profiles were compared between the two experimental groups, marked differences were noted between *L. crispatus* infected organoids and those infected by *F. vaginae* (Figure 4). Among the most highly differentially expressed genes, transcripts from the CXCL family, including CXCL1,2,3,5,6,10, were highly enriched in *F. vaginae* infected organoids, which may contribute to leukocyte recruitment. Several other genes, such as TGFB1 or TRADD, are upregulated by *L. crispatus* (Figure 4).

**Figure 3.**
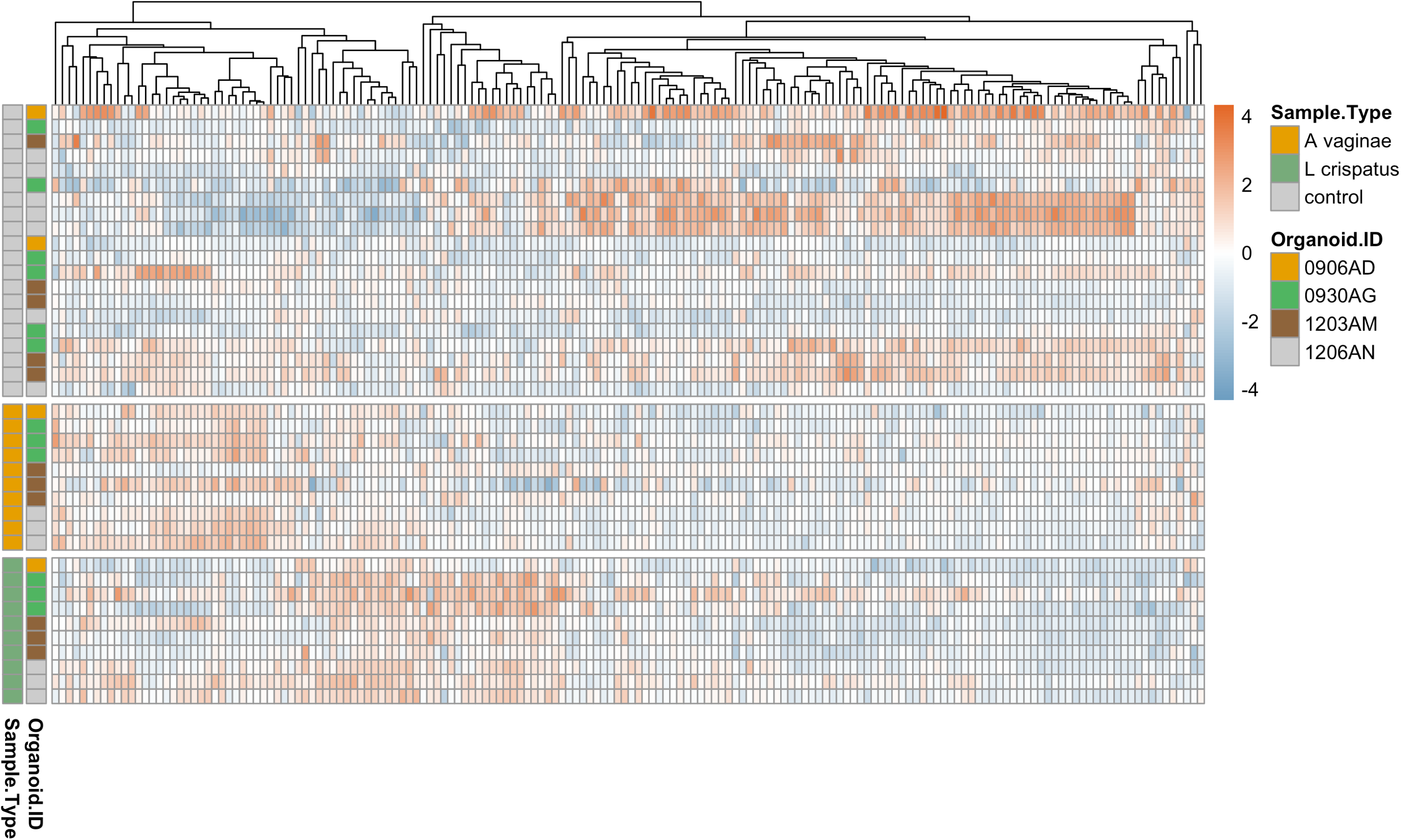
Heatmap overview of gene expression levels of all samples as grouped by treatment groups. Top panel (grey left bar): negative controls without bacterial co-culture. Middle panel (orange left bar): organoids co-cultured with *L. crispatus*. Bottle panel (green left bar): organoids co-cultured with *F. vaginae*. Individuals are color coded in second left bar (organoid ID). Each row is a sample. Each column is a gene. Heatmap scale: −4 to 4 (blue to orange) indicates low to high level of gene expression.

**Figure 4.**
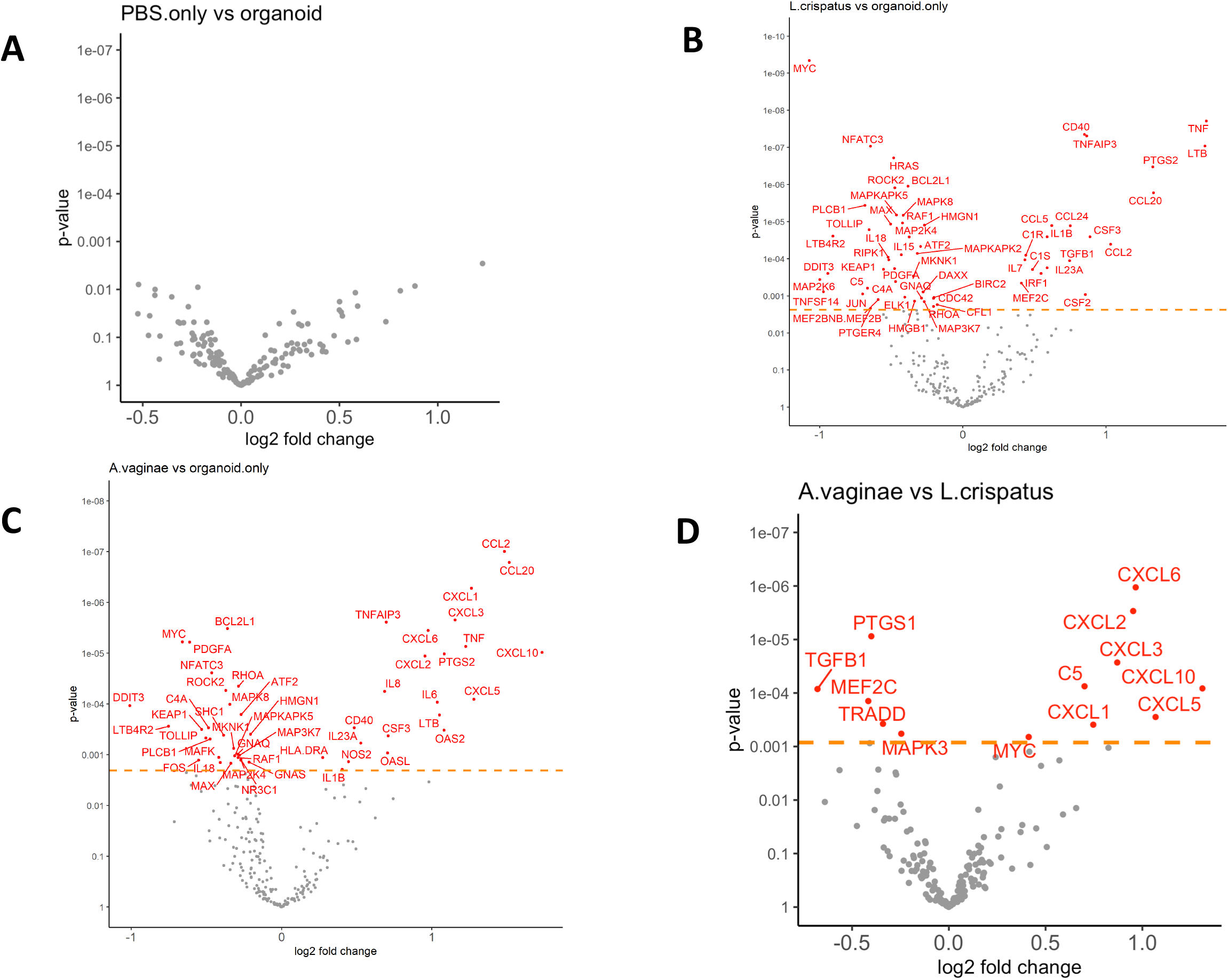
Differential gene expressions between treatment groups as shown in volcano plots. A. No significant gene expression changes caused by adding PBS in the culture media. B-C. Bacterial co-culture (B. *L. crispatus*, C. *F. vaginae*) resulted in multiple gene expression changes compared to negative controls. D. Differentially expressed genes (in red) between the two bacterial culture groups.

With flow cytometry, we assessed the presence of immune cells during the FT organoid culturing period. On the day of tissue collection, 22% of cells were CD45+ immune cells. On the next day after 24 hours of culture in 2D growth media, only 11% of cells were immune cells. The CD45+ immune cells disappeared rapidly within the first five days of 3D culture to ≤1% and were undetectable at later timepoints (Day 6-9 of 3D culture, <1% of cells were CD45+, data not shown) (Figure 5). The lack of immune cells in mature organoids (on average 14 days in 3D culture) indicated the inflammatory response observed with bacterial culture was generated by the epithelial cells in the organoids.

**Figure 5.**
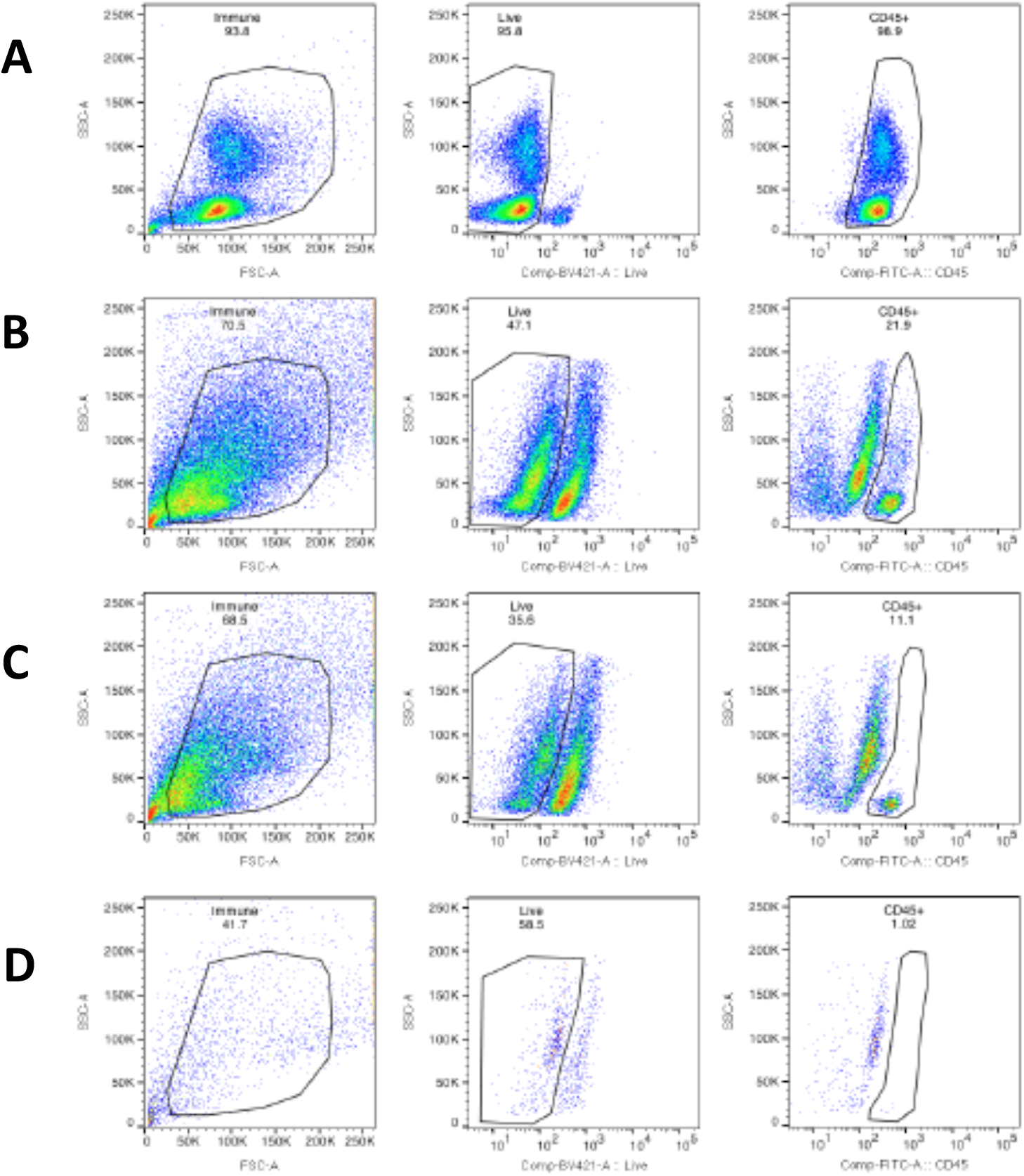
Disappearance of immune cells in FT organoid culture. The percentage of viable CD45+ immune cells were analyzed by flow cytometry at various time points during the organoid culture. Gating strategy for each sample; left panel: gating of single cells by forward and side scatter, center panel: gating of live cells (Live/DEAD negative), right panel: gating of CD45+ cells. A. Positive control using PBMCs. B. Day 0 of FT tissue collection, 22% of cells were immune cells. C. Day 1 of 3D organoid culture, 11% of cells were immune cells. D. Day 5 of 3D organoid culture, ≤1% of immune cells remaining.

## DISCUSSION

By inoculating the culture media with two common vaginal bacterial species, we were able to elicit acute inflammatory responses from the epithelial cells in the FT organoids. Our results indicate the divergence in the inflammatory responses provoked by pathogenic and non-pathogenic vaginal bacterial species. We demonstrate rapid disappearance of immune cells in the FT organoid culture, so this inflammatory response was generated directly by the FT epithelial cells.

Among the genes upregulated in *F. vaginae* infected organoids in comparison to the negative controls and to the *L. crispatus* infected organoids, multiple genes encoding C-X-C motif chemokine ligand (CXCL) family were represented. The CXCL family widely participates in immune cell recruitment, specifically, attracts monocytes and neutrophils, and is involved in the pathogenesis of inflammatory diseases such as endometriosis and its associated infertility^27–29^ as well as angiogenesis and tumor progression in ovarian and cervical cancer^30,31^. C5 that is highly upregulated in the *F. vaginae* infected organoids is a chemotactic molecule that directly attracts neutrophils, eosinophils, basophils, and monocytes to the sites where complement is activating^32^. Therefore, it is not surprising for the FT organoid epithelial cells to increase the production of these chemokines in reaction to pathogenic bacteria such as *F. vaginae*.

Multiple members of the CC-chemokine ligand (CCL) family were upregulated in both *F. vaginae* and *L. crispatus* infected organoids. These chemokines attract monocytes, T cells and eosinophils to inflammatory sites. In a previous publication, when these bacterial species were incubated with vaginal epithelial cell aggregrates, it was found that *F. vaginae* led to a strong cytokine response including upregulation of CCL20 as seen in our study with FT organoids; however, *L. crispatus* at a lower concentration elicited much lower and in some cases unmeasurable cytokine release from the vaginal epithelia cells^33^. Even though *L. crispatus* is a non-pathogenic bacteria in the vagina, it does not normally reside in the FT in high concentration, which may explain why *L. crispatus* elicited inflammatory response when incubated with the FT organoids in our study, but minimal repsonse when incubated in low concentration with vaginal epithelial cells in previous study.

Interestingly, TGFB1, the gene that encodes TGF-β1, is upregulated by *L. crispatus*, but not by *F. vaginae*, and TGF-β1 plays an important role in the maintenance of immune homeostasis and self-tolerance by inducing unresponsiveness in naïve T cells and preventing lymphocyte activation^34^. This may indicate a potential mechanism that *L. crispatus* utilizes to promote the tolerance of normal microbiome in the female genital tract. Another interesting finding was the downregulation of MAPK pathway genes in *L. crispatus* infected organoids, as MAPK genes were found to be some of the key genes in novel pathways identified in PID associated infertility^35^.

To our knowledge, no previous publication has studied the inflammatory response of FT organoids to vaginal bacteria. Our study indicates that FT organoids can be a useful model system to study the host-pathogen interaction during bacterial infection which may facilitate mechanistic investigations in PID and its contribution to tubal factor infertility and ovarian carcinogensis. Some of the technical strengths of this study include: three different types of negative controls were used; the replicates showed little intra-individual variations; high RNA integrity was observed across the samples (RIN > 9); all samples were processed by the same individuals and analyzed on the same Nanostring panels at the same time, thus minimizing batch effects. Inter-individual variations were noted in the inflammatory response to either bacteria; while commonly observed in other organoids and used as a tool for precision medicine^36–38^, a much larger sample size would be required to confirm whether consistent individual heterogeneity exists in FT organoid inflammatory responses. Building on the feasibility established in this pilot study, we are developing a more complex FT organoid culture system to better investigate the host-pathogen interaction, such as incorporating the immune cells into the culture, co-culturing multiple bacterial species together to better represent the normal or pathological genital tract microbiome, and evaluating the genome-wide changes in gene expression rather than focusing on the inflammatory genes only. These efforts will lead to a better understanding of the local microenvironmental changes during acute and chronic infections which may lead to infertility or cancer initiation.

## CONCLUSIONS

In this pilot study, we demondate that epithelial cells in patient tissue derived FT organoids respond to acute bacterial infection with upregulation of inflammatory genes specific to each vaginal bacterial species in culture: non-pathogenic *L. crispatus* or pathogenic *F. vaginae*. FT organoids is a useful model system to study the host-pathogen interaction during bacterial infection which will facilitate mechanistic investigations in PID and its contribution to tubal factor infertility and ovarian carcinogensis. Further studies are ongoing to develop a more sophisticated FT organoid culture system to better mimic the FT microenvironmental changes during acute and chronic infections leading to infertility or cancer initiation.

## Acknowledgements

The study was supported by *Akiko Yamazaki and Jerry Yang* Faculty Scholar Fund in Pediatric Translational Medicine, Stanford Maternal and Child Health Research Institute, and the Dunlevie Maternal-Fetal Medicine Center for Discovery, Innovation and Clinical Impact at Stanford University (to BY).

## Funding information

The study was funded by *Akiko Yamazaki and Jerry Yang* Faculty Scholar Fund in Pediatric Translational Medicine, Stanford Maternal and Child Health Research Institute (to BY), NIH K08CA222835 (to BY), NIH R01AI139189 (to CH, DNF, BY).

## Conflict of interest disclosures

The authors declare that there are no conflicts of interests.

## Authors contribution statement

BY conceived and designed the study, drafted the manuscript. BY, EBN, SM, SP, SS, DV, CL collected and analyzed the data. BY, CH, DNF obtained funding and edited the manuscript. All authors have read and approved the final manuscript.

## Notes

**Funding Statement**: The study was funded by *Akiko Yamazaki and Jerry Yang* Faculty Scholar Fund in Pediatric Translational Medicine, the Stanford Maternal and Child Health Research Institute (to BY), NIH K08CA222835 (to BY), NIH R01AI139189 (to CH, DNF, BY).

### Competing Interest Statement

The authors have declared no competing interest.

